# Gene Expression-Based Drug Repurposing to Target Ageing

**DOI:** 10.1101/253344

**Authors:** Handan Melike Dönertaş, Matías Fuentealba Valenzuela, Linda Partridge, Janet M. Thornton

## Abstract

Ageing is the largest risk factor for a variety of non-communicable diseases. Model organism studies have shown that genetic and chemical perturbations can extend both life- and health-span. Ageing is a complex process, with parallel and interacting mechanisms contributing to its aetiology, posing a challenge for the discovery of new pharmacological candidates to ameliorate its effects. In this study, instead of a target-centric approach, we adopt a systems level drug repurposing methodology to discover drugs that could combat ageing in human brain. Using multiple gene expression datasets from brain tissue, taken from patients of different ages, we first identified the expression changes that characterise ageing. Then, we compared these changes in gene expression with drug perturbed expression profiles in the Connectivity Map. We thus identified 24 drugs with significantly associated changes. Some of these drugs may function as anti-ageing drugs by reversing the detrimental changes that occur during ageing, others by mimicking the cellular defense mechanisms. The drugs that we identified included significant number of already identified pro-longevity drugs, indicating that the method can discover de novo drugs that meliorate ageing. The approach has the advantages that, by using data from human brain ageing data it focuses on processes relevant in human ageing and that it is unbiased, making it possible to discover new targets for ageing studies.

## Introduction

Life expectancy has increased steadily in many countries worldwide. Since ageing is the major risk factor for multiple pathologies, including cardiovascular diseases, neurodegenerative disorders, and cancer (Niccoli & Partridge, 2012), finding interventions that can increase health during ageing is of importance. Lifespan of laboratory model organisms can be greatly extended by genetic and environmental interventions, which also improve health and function during ageing (Clancy et al., 2001; Lucanic, Lithgow, & Alavez, 2013; Xiao et al., 2013). Many of these interventions target components of the nutrient-sensing network, and decrease the activity of IGF/Insulin and/or TOR signalling (Fontana, Partridge, & Longo, 2010). Moreover, dietary restriction (DR), decreased food intake without malnutrition, can increase lifespan, and further supports the importance of nutrient sensing pathways in ageing (Fontana & Partridge, 2015).

Pharmacological intervention can also extend animal lifespan. The DrugAge database reports drug-induced lifespan extensions up to 1.5-fold for *C. elegans*, 1.1-fold for *D. melanogaster*, and 31% for *M. musculus* (Barardo et al., 2017). Some of these chemicals may mimic the effects of DR (Fontana et al., 2010). For example, resveratrol, which induces a similar gene expression profile to dietary restriction (Pearson et al., 2008), can increase lifespan of mice on a high-calorie diet, although not in mice on a standard diet (Strong et al., 2013). Rapamycin, directly targets the mTORC1 complex, which plays a central role in nutrient sensing network and has an important role in lifespan extension by DR (Mair & Dillin, 2008). Rapamycin extends lifespan by affecting autophagy and the activity of the S6 kinase in flies. However, it can further extend the fly lifespan beyond the maximum achieved by DR, suggesting that different mechanisms might be involved (Bjedov et al., 2010). Nevertheless, the mechanisms of action for most of the drugs are not well known.

Several studies have taken a bioinformatics approach to discover drugs that could extend lifespan in model organisms. For instance, the Connectivity Map, a database of drug-induced gene expression profiles, has been used to identify DR mimetics, and found 11 drugs that induced expression profiles significantly similar to those induced by DR in rats and rhesus monkeys (Calvert et al., 2016). Another study generated a combined score reflecting both the ageing relevance of drugs based on the GenAge database and GO annotations as well as the likely efficacy of the drugs in model organisms, using structural analyses and other criteria such as solubility (Ziehm et al., 2017). A machine learning approach has been used to identify pro-longevity drugs based on the chemical descriptors of the drugs in DrugAge database and GO annotations of their targets (Barardo et al., 2017). By using DrugAge as a training set, the results reflect the known pathways in ageing, and thus identified anti-cancer and anti-inflammatory drugs, compounds related to mitochondrial process and gonadotropin-releasing hormone antagonists. Another study took a pharmacological network approach to characterise anti-ageing drugs, first screening a large library of 1280 compounds for lifespan extension in *C. elegans*. The 60 hits from the screen were used to construct a pharmacological network, and clustered in certain pharmacological classes, mainly related to oxidative stress (Ye, Linton, Schork, Buck, & Petrascheck, 2014).

While most studies have focussed on model organisms, one study used the known pro-longevity drugs from the Geroprotectors database (Moskalev et al., 2015) and asked if these could be functional in humans (Aliper et al., 2016). Using young and old human stem cell expression profiles, they calculate a geroprotective score based on the GeroScope algorithm, which scores drugs based on the drug targets and age-associated expression changes in related pathways (Zhavoronkov, Buzdin, Garazha, Borisov, & Moskalev, 2014). Testing the top hits in senescent human fibroblast cultures, they suggest several geroprotectors for humans as well as showing the potential in using human gene expression data for drug studies.

Here, we extended the approaches to identification of new anti-ageing drugs, by focusing directly on human ageing. We used a framework that does not require any prior knowledge and is thus robust to biases in the literature and databases on ageing. Through a meta-analysis of multiple gene expression datasets, we first compiled a robust signature that characterises ageing in human brain. We then used drug-induced RNA expression profiles deposited in the Connectivity Map (CMap) (Lamb, 2006) to identify a list of potential drug candidates that could influence human brain ageing. We then assessed the performance of the method in relation to previous knowledge and identified novel candidate geropotective drugs.

## Results

### Analysis of age-related changes in RNA expression in human brains

We analysed data from seven, published, microarray-based studies of age-related changes in RNA expression (Barnes et al., 2011; Berchtold et al., 2008; Colantuoni et al., 2011; Kang et al., 2011; Lu et al., 2004; Maycox et al., 2009; Somel et al., 2010, 2011). The data came from 22 different brain regions, and the ages of the donors ranged from 20 to 106 years (Figure 1a, Figure S1). The data for each brain region in each study were analysed separately, resulting in 26 datasets.

**Figure 1:**
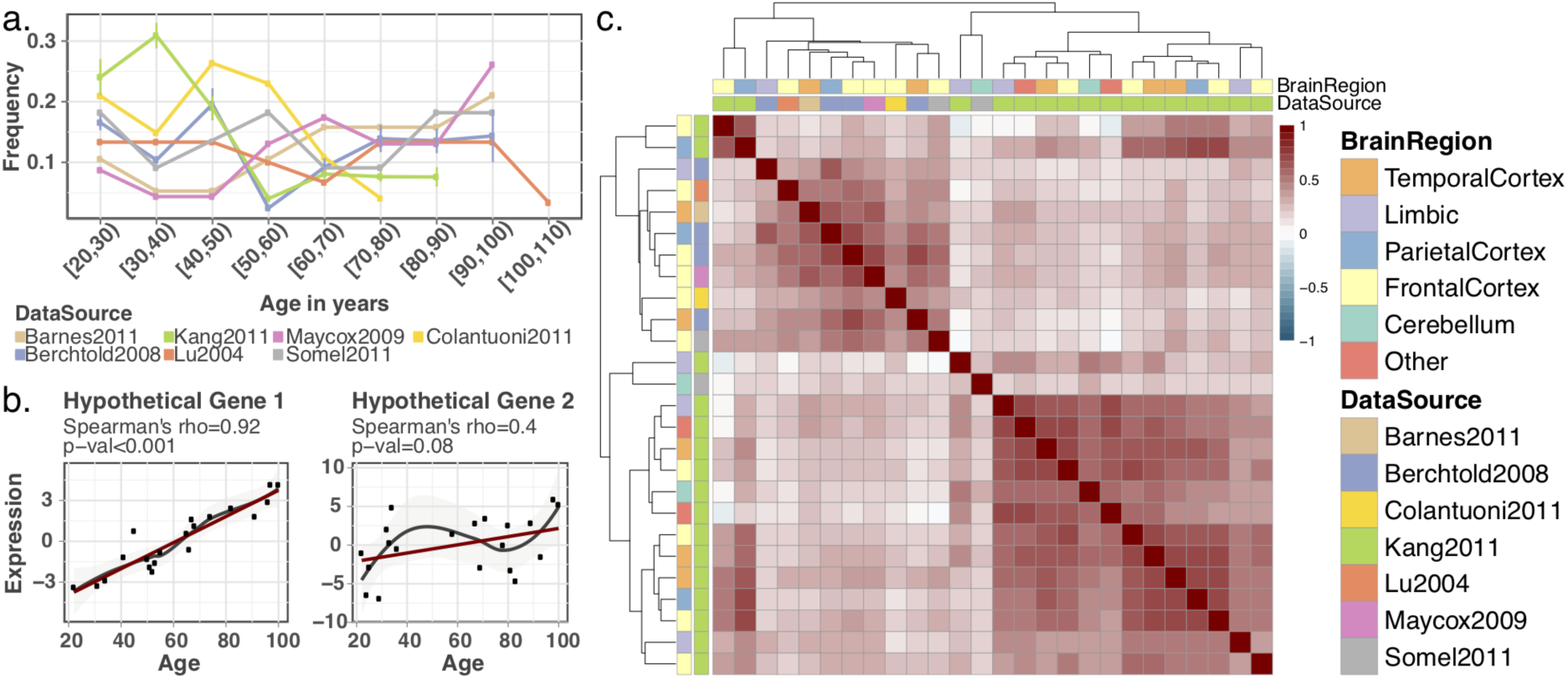
a) Age distribution of the brains from which the datasets used in the study were derived. The error bars show the standard deviation of the sample frequency for different brain regions in data sources with multiple brain regions. b) Hypothetical gene expression plots, demonstrating how Spearman’s correlation coefficient and p-value behave when the association is weak or non-monotonic. c) Pairwise Spearman’s rank correlation coefficients across datasets. The intensity of the colours on the heatmap shows the magnitude of the correlation coefficient.

To characterise the association between the gene expression and age, we calculated the Spearman’s correlation between the expression level and age, for each gene, in each dataset separately. We first calculated the number of significant changes (FDR corrected p<0.05) in each dataset (Figure S2). While there were two datasets with a large number of significant changes, most of the datasets did not show substantial significant change. This can be explained by several factors, most importantly i) most of the datasets had a small sample size, providing insufficient power to detect changes in most of the cases, and ii) Spearman’s correlation test calculates significant monotonic changes, whereas it is likely that many of the changes are not exclusively monotonic throughout ageing. Thus, we applied another approach, using the correlation coefficient to capture significant trends across datasets, instead of within a dataset (see Methods). While the p-value is affected by the number of the samples and the strength of the monotonic relationship (Figure 1b), the sign of the correlation coefficient can be used to capture consistent trends of up- or down-regulation once coupled with an appropriate testing scheme. This strategy requires the datasets to be concordant and reflect genuine age-related changes. We first investigated if this assumption was valid. To assess the concordance among datasets, we used Spearman’s correlation coefficients and calculated the correlation between expression-age correlations between datasets (Figure 1c). We observed a weak correlation with a median pairwise correlation coefficient of 0.29. To calculate the significance of this correlation, we developed a stringent permutation scheme specifically designed to account for the dependence between genes as well as the datasets (see Methods for detail). We concluded that a median correlation coefficient of 0.09 would be expected by chance and that our observation (median rho=0.29), is statistically significant (p<0.001). Based on these correlations, datasets clustered according to the data source rather than to the brain region. This observation is in line with the previous studies suggesting that ageing-related changes are small and heterogeneous, making them difficult to detect (Somel, Khaitovich, Bahn, Pääbo, & Lachmann, 2006). We therefore tested for significant correlations across datasets from different studies. When we excluded the correlation coefficients among the datasets generated by the same studies, we still observed a significant correlation coefficient of 0.22 (permutation test p<0.001, rho=-0.002 would be expected by chance), showing that we have significant correlations among different data sources as well. Using these correlations, we proceeded to compile the ageing-signatures, reflecting consistent trends.

### Defining the ageing signature

To construct a robust ageing signature, we identified the age-related changes that were observed across all datasets, irrespective of the effect size. We thus focussed on global age-related changes in the brain, rather than region-specific changes, and the set of genes that showed gene expression changes in the same direction across all datasets (Figure 2a). This profile consisted of only 100 up- and 117 down-regulated genes (Table S2, Figure S3-4), ’the ageing signature’.

**Figure 2:**
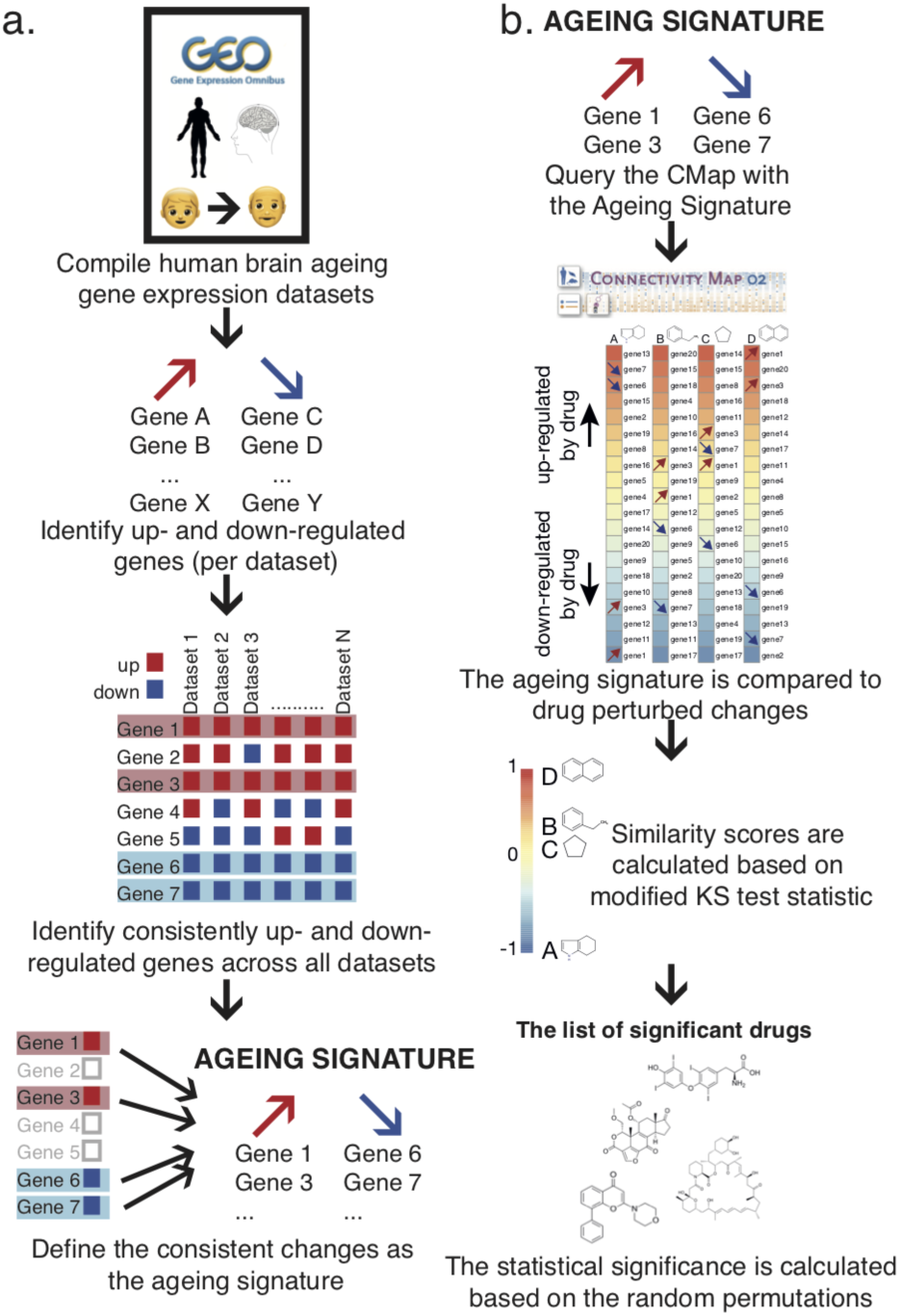
Method summary for a) compiling the ageing signature and b) the CMap algorithm.

To establish the robustness of the ageing signature, we calculated the statistical significance of the number of consistent changes with the same permutation scheme used to test the correlations among datasets. This methodology randomizes the age of each individual, making it possible to test the null hypothesis where there is no association between expression and age while retaining the dependence between genes and datasets (see Methods for details). The number of consistent expression changes across brain regions was significant (p<0.001, Figure S6a-b), establishing that the ageing signature indeed has biological meaning.

To further test the robustness of the ageing signature, we used an independent data set, consisting of gene expression in human brain generated by the GTEx Consortium (Ardlie et al., 2015), consisting of data from 99 individuals, 13 brain regions and ages between 20-79 (Figure S1, Table S1). These data were generated using RNA-Seq, allowing us to assess the robustness of the ageing signature to different technology platforms. We used pipeline previously applied to the microarray data to calculated age-related expression changes for each gene in each brain region separately. The pairwise correlations between the GTEx datasets were higher than with the other dataset, and they tended to cluster together (Figure S5). We found 1189 up- and 1352 down-regulated genes that showed the same direction of change across all GTEx brain regions (Table S2), compared with only 100 and 117 in the microarray ageing signature. A likely explanation is that samples from different brain regions from the same individuals were used in GTEx, while the microarray ageing signature combined seven independent studies and different microarray platforms. The numbers of shared expression changes based on permutations were 127 and 131.5, for down- and upregulated genes, suggesting a higher false positive rate in the GTEx dataset. Nevertheless, the numbers of consistent up- and down-regulated genes in the GTEx dataset were also significant (p=0.001, Figure S6c-d). The numbers of common up- and down-regulated genes across the GTEx and microarray signatures were 50 and 48, respectively, both statistically significant (binomial test p < 2.2e-16 for both), demonstrating that the ageing signature was reproducible.

### Biological processes associated with the ageing signature

We next investigated the biological processes associated with the microarray ageing signature. Using the genes that were consistently expressed in all data sources as background, we did Gene Ontology enrichment tests for consistently up- and down-regulated genes, separately (Figure 3, Table S3 (up-regulated), Table S4 (down-regulated)). Down-regulated genes were enriched in synaptic functions and biosynthetic processes (FDR corrected p<0.05), while differentiation and proliferation-related categories showed enrichment for the up-regulated genes (FDR corrected p<0.05). These results are consistent with the findings of earlier brain ageing transcriptome studies (Lu et al., 2004; Naumova et al., 2012; Xue et al., 2007). Oddly, ossification-related biological processes also showed significant enrichment for the up-regulated genes. However, except for one gene, these ossification-related categories shared all genes with the more generic development-related categories. Thus, this result could be interpreted as a general up-regulation of the development-related processes rather than ossification-related categories.

**Figure 3:**
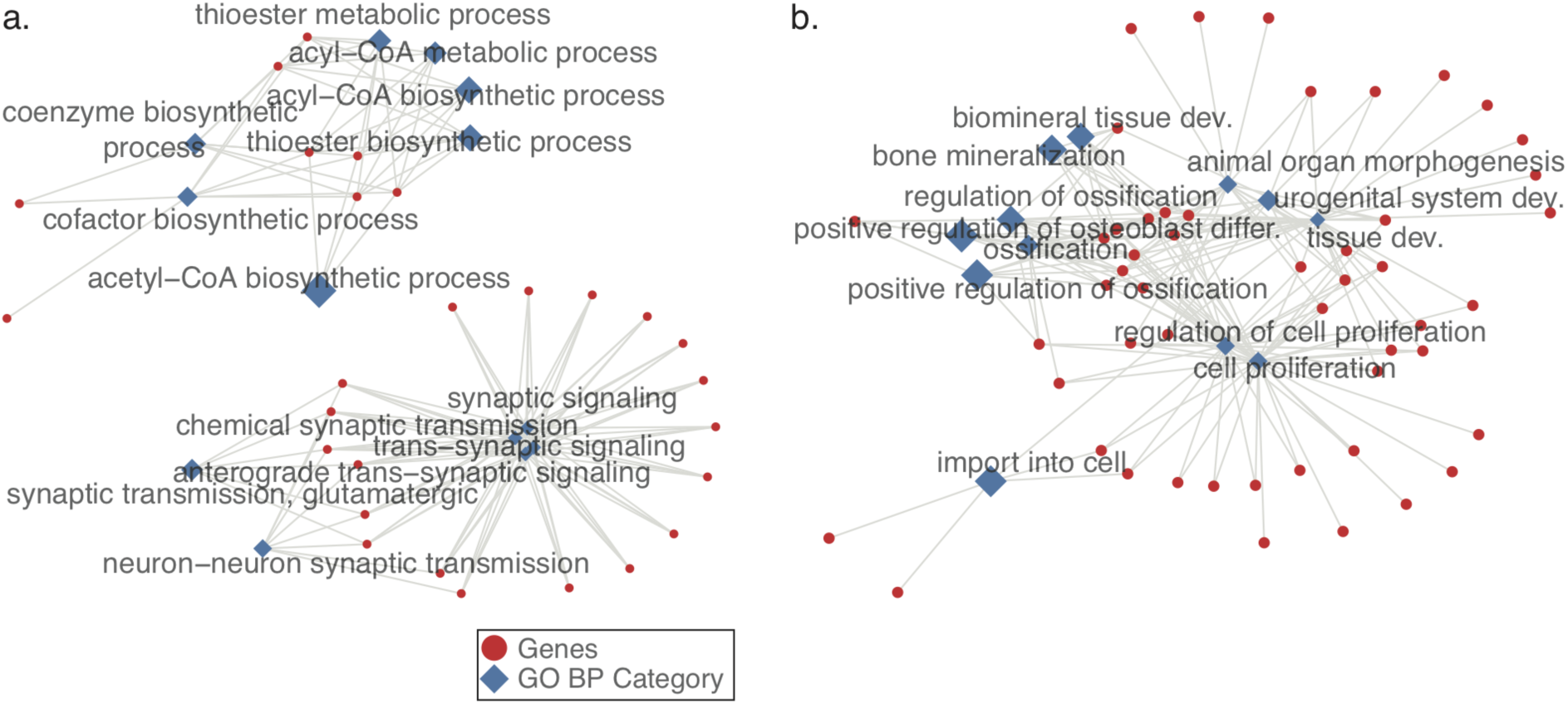
Gene Ontology Biological Process Categories significantly enriched in a) down- and b) up-regulated genes in the microarray ageing signature. Red circles represent the genes, diamonds show the significantly associated GO Categories, where FDR adjusted p<0.05. The size of the diamonds represents the effect size (odds ratio).

We repeated the enrichment analysis using the GTEx ageing signature and found 194 and 256 GO BP categories as significantly associated with down- and up-regulated genes, respectively (TableS7-8). Since the number of genes in the GTEx signature is higher, we had more power to detect smaller changes and thus had a higher number of significant associations. However, the effect sizes (odds ratios) for each GO BP category calculated for microarray and the GTEx ageing signature were correlated (FigureS7). Correlations between the odds ratios calculated for all of the GO categories calculated in both methods were 0.46 and 0.37, for the enrichment in the down- and up-regulated genes, respectively. Correlations increase when we considered only the GO categories that are significantly associated with at least one of the ageing signatures; 0.55 and 0.60, for the enrichment in the down- and up-regulated genes, respectively. This further shows that the ageing signatures are robust. The categories enriched in down-regulated genes included biological processes related to neuronal and synaptic functions, autophagy, post-translational modifications, and translation (see Table S7 for the full list). Processes related to response pathways, immune response, macromolecule organisation and lipid metabolism showed enrichment in up-regulated genes (see TableS8 for the full list). Interestingly, categories related to ossification were also among the GO categories significantly associated with up-regulation, based on GTEx data.

### Mapping the ageing signature onto drug-perturbed expression profiles

The Connectivity Map (CMap) is a database of drug perturbed gene expression profiles (Lamb, 2006). It consists of 6100 gene expression profiles for 1309 drug perturbation experiments performed on five different cell lines. The CMap algorithm uses a modified Kolmogorov-Smirnov test statistic to calculate the similarity of a drug-perturbed expression profile to the gene expression profile used to query the database. A positive similarity score means that the drug-perturbed expression profile is similar to the query, whereas a negative score indicates a negative correlation (Figure 2b). Based on the random permutations, the statistical significance of the similarity score for each drug is calculated. Thus, the p-value shows the probability of finding the same association when a random signature is supplied. We queried the CMap database and identified drugs that showed significant associations in either direction with the ageing signatures. To determine the robustness of this procedure, we queried the CMap data using the microarray ageing signature, and the top 500 up- and 500 down-regulated genes from the GTEx ageing signature (see Methods). The correlation was significant (r=0.52, p<2.2e-16, Figure S3a) showing that the two ageing signatures produce reproducible overlaps with the CMap database.

Querying the CMap database, we identified 13 drugs significantly associated (FDR corrected p<0.05) with the microarray ageing signature (Table1, Figure 4). Four of these drugs were previously shown to extend lifespan in worms or flies in at least one experiment (Table S9). The number of pro-longevity drugs re-discovered using this methodology was statistically significant (p=0.004), and only one drug would be expected based on 10,000 random permutations of drugs. Repeating the same analysis with the GTEx ageing signature, we identified 18 drugs, seven of which were in common with the microarray results, including the four known pro-longevity drugs. In total, 24 drugs were significantly associated with at least one of the ageing signatures. The correlation between the drug similarity scores for these 24 drugs calculated based on the microarray and GTEx data was 0.88 (p<9.44e-09, Figure S3b), indicating high concordance. Since the similarity scores show high correlation, the rest of the results will be presented for the 24 drugs that are associated with at least one of the ageing signatures.

**Figure 4:**
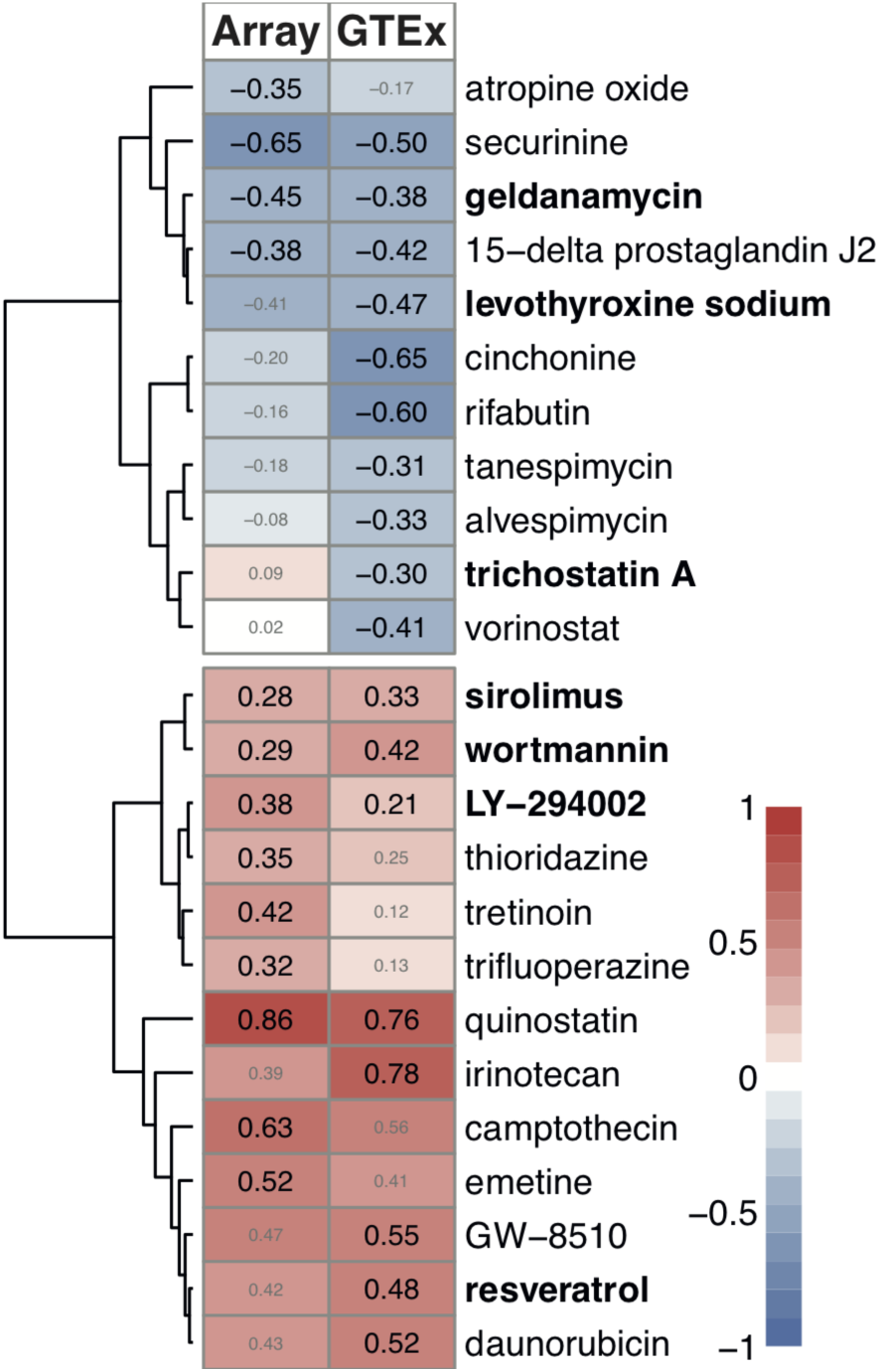
Similarity score table for the drugs having at least one significant association to the ageing signatures. Each row corresponds to a drug and columns correspond to two independent ageing signatures – using the microarray and the GTEx datasets. The size of score labels indicates the significance of the results (FDR corrected p<0.05). The row labels written in bold indicates the drugs in the DrugAge database.

Overall, the method re-discovered seven known pro-longevity drugs in DrugAge database (p=0.00023, based on 100,000 random permutations); resveratrol, LY-294002, wortmannin, sirolimus (also known as rapamycin), trichostatin A, levothyroxine sodium, and geldanamycin (Table S9).

### Targets of the drugs

Next, we investigated the targets of these 24 drugs, using the ChEMBL, PubChem and DrugBank databases as well as through manual curation of the literature (Table 1), and whether these targets were previously implicated in ageing, using GenAge human and model organism databases (Figure 5). Except for four (rifabutin, securinine, thioridazine, trifluoperazine); all drugs or their target genes had been previously implicated in ageing. Moreover, the drug-target association network showed several clusters with multiple drugs sharing the same targets: i) quinostatin was in the same cluster with two known pro-longevity drugs, wortmannin and LY-294002, targeting PI3K subunits, ii) tanespimycin and alvespimycin shared the same target with another DrugAge drug, geldanamycin, targeting HSP90, iii) vorinostat shared one of its targets, HDAC6, with trichostatin A, another DrugAge drug, iv) thioridazine and trifluoperazine had dopamine and serotonin receptors as targets and v) irinotecan and camptothecin shared TOP1 as their target. The fact that drugs targeting the same proteins / acting through the same mechanism had similar CMap similarity scores (Figure 4) further shows that our results are biologically relevant and reflects potential mechanisms to target ageing.

**Table 1:** The drugs that are significantly associated (FDR corrected p<0.05) with at least one of the ageing signatures. Drug names in bold shows the drugs in DrugAge database. ‘Score’ is the mean similarity score given in the CMap output, based on KS test. The similarity scores denoted with (*) show the significant associations. The list is ordered by the mean of the similarity scores from negative to positive. Target or mechanism of action is manually curated from literature (the relevant literature is given in the SI) or extracted from CHEMBL, DrugBank, and PubChem databases. The targets written in bold are found in the GenAge model organism or GenAge human databases.

| DRUG NAME | ARRAY SCORE | GTEX SCORE | TARGET OR MECHANISM OF ACTION |
| --- | --- | --- | --- |
| Securinine | -0.65 (*) | -0.50 (*) | GABRA1-5, GABRB1-3 |
| **Levothyroxine sodium** | -0.41 | -0.47 (*) | THRA, **THRB** |
| Cinchonine | -0.2 | -0.65 (*) | **CYP2D6** |
| **Geldanamycin** | -0.45 (*) | -0.38 (*) | **HSP90AA1** |
| 15-delta prostaglandin J2 | -0.38 (*) | -0.42 (*) | **PPARG** |
| Rifabutin | -0.16 | -0.6 (*) | BCL6 |
| Atropine oxide | -0.35 (*) | -0.17 | - |
| Tanespimycin | -0.18 | -0.31 (*) | **HSP90AA1** |
| Alvespimycin | -0.08 | -0.33 (*) | **HSP90AA1** |
| Vorinostat | 0.02 | -0.41 (*) | **HDAC1, HDAC2, HDAC3, HDAC6** |
| **Trichostatin A** | 0.09 | -0.3 (*) | HDAC6, HDAC7, HDAC8 |
| Trifluoperazine | 0.32 (*) | 0.13 | DRD2, DRD3, DRD4, HTR2A, HTR2C |
| Tretinoin | 0.42 (*) | 0.12 | **RARA, RARB, RARC** |
| **LY-294002** | 0.38 (*) | 0.21 (*) | **PI3KCG** |
| Thioridazine | 0.35 (*) | 0.25 | DRD2, DRD3, DRD4, HTR2A, HTR2C |
| **Sirolimus** | 0.28 (*) | 0.33 (*) | **mTOR** |
| **Wortmannin** | 0.29 (*) | 0.42 (*) | **PI3KR1, PI3KCA, PI3KCG** |
| **Resveratrol** | 0.42 | 0.48 (*) | SULT1B1, YARS, LTA4H, TTR, NQO2, **PTGS2**, PTGS1, MAT2B, CSNK2A1, **CYP3A4, ESR1, PPARG, SIRT1, SIRT5, CYP1A2, CYP1A1, CYP1B1, NCOA2, TNNC1** |
| Emetine | 0.52 (*) | 0.41 | Protein Synthesis Inhibition |
| Daunorubicin | 0.43 | 0.52 (*) | **TOP2A, TOP2B** |
| GW-8510 | 0.47 | 0.55 (*) | CDK2, **CDK5** |
| Irinotecan | 0.39 | 0.78 (*) | **TOP1** |
| Camptothecin | 0.63 (*) | 0.56 | **TOP1** |
| Quinostatin | 0.86 (*) | 0.76 (*) | **PI3KCA** |

**Figure 5:**
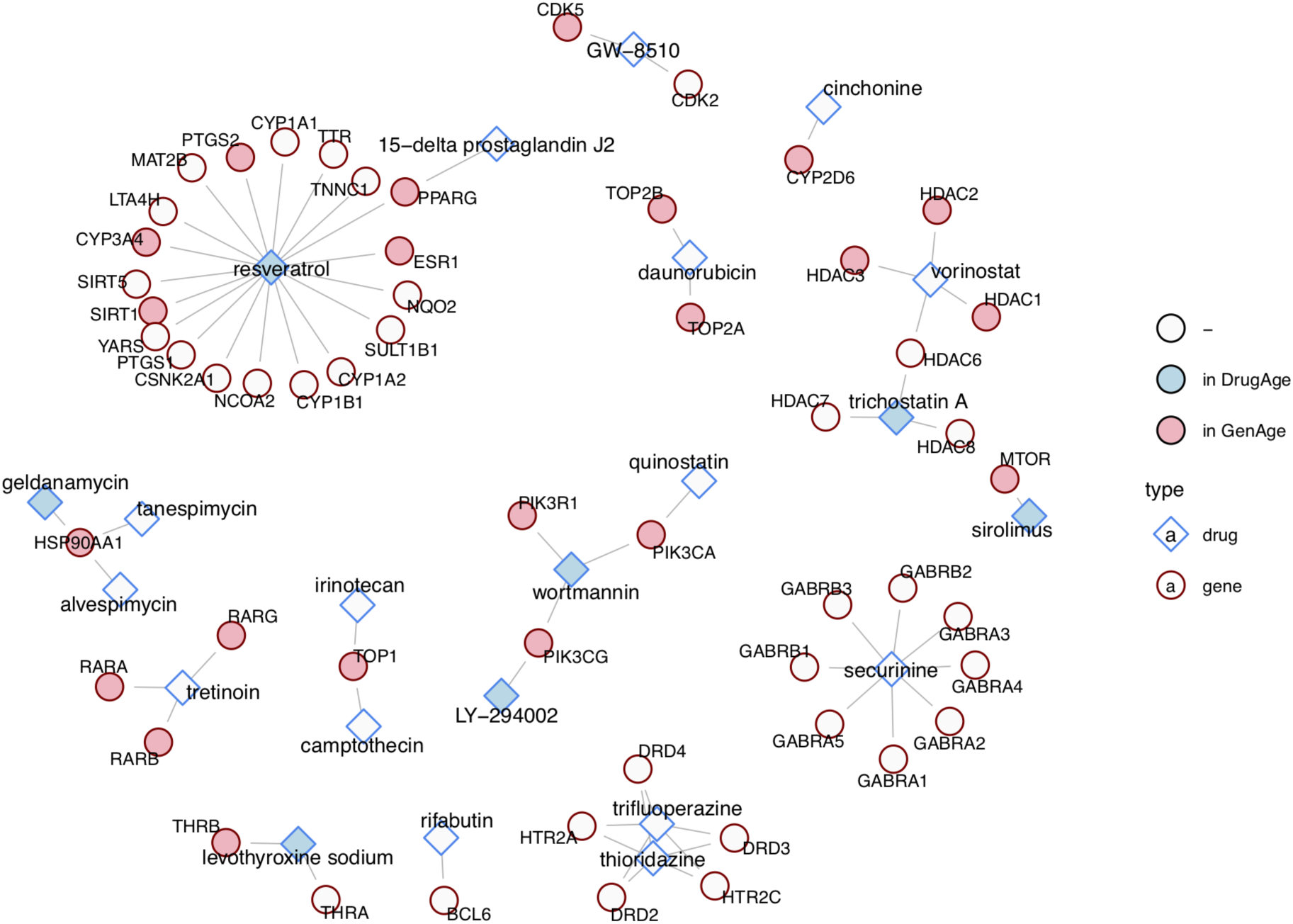
Schematic representation of the drug-target associations as a network. Blue and red nodes show drugs and targets, respectively. The drugs with a light blue background are present in DrugAge database and the targets with a pink background are in either GenAge model organism or GenAge human databases.

### Drugs can act both by reversing ageing effects and mimicking responses

The general expectation from an ‘omics-based drug repurposing study is the identification of drugs that can reverse the abnormalities detected in the disease state i.e. identification of drugs with negative similarity scores (Duran-Frigola, Mateo, & Aloy, 2017). Following the same logic, one might expect drugs with anti-ageing potential to have negative scores. Interestingly, some of the known pro-longevity drugs had positive similarity scores to the ageing signatures, meaning that the drug-induced profile was similar to the ageing signature. A plausible explanation for this observation is that ageing signatures may partly reflect cellular defense responses, helping to alleviate the damaging effects of ageing.

### Characterising the biological functions associated with pro-longevity drugs

In order to identify the biological processes associated with the changes that were reversed or mimicked by the pro-longevity drugs, we used the drugs documented in DrugAge, that were re-discovered in our analysis. We grouped the microarray ageing signature into five categories, based on the expression changes in ageing (up or down), and the pro-longevity drug-induced profile (up, down or inconsistent) (TableS5). To compile the pro-longevity drug profile, for each probe-set in the microarray ageing signature, we asked if the seven DrugAge drugs induced similar changes. If the same direction of change was induced by more than half of these DrugAge drugs, then we included these changes in the pro-longevity drug profile (see Methods for the details). We then analysed the biological processes associated with the genes in these categories. The number of genes is small, with no significant changes after multiple test correction. We therefore report the associations based on the highest odds ratios only. For genes down-regulated in ageing, the changes mimicked by the drugs were associated with autophagy and metabolic processes (Table S6), while for up-regulated genes, pro-longevity drugs tended to mimic the changes in protein complex / cellular complex assembly-related functions and to reverse the changes observed in protein localisation and immune-related functions (TableS6). These findings are consistent with the mechanism of action for the most well-known pro-longevity drugs. For example, sirolimus (rapamycin) is an immunosuppressant approved for human use, and similar drugs can enhance the response of elderly humans to immunization against influenza (Mannick et al., 2014).

### Similarity among significant drugs based on the expression changes at the functional level

In order to analyse the similarities among drugs based on expression level changes, we performed a gene-set enrichment analysis (GSEA) for the drug-induced expression profiles, including all genes irrespective of whether a given gene is in the ageing signature (see Methods). To measure the similarity between drugs, we calculate the Spearman’s rank correlation coefficients between the enrichment scores and then cluster drugs based on these correlation coefficients. Notably, drugs targeting the same proteins or pathways, e.g. PI3K inhibitors LY294002, wortmannin and quinostatin, clustered together. Using this functional level approach, we grouped drugs into four groups: i) known pro-longevity drugs, ii) drugs clustering together with at least one pro-longevity drug, iii) drugs which clustered together but did not cluster with any known pro-longevity drugs, and iv) drugs which did not cluster with any other drugs (Figure S10).

### Ageing signature in other tissues

Since our analysis is based on an ageing signature compiled using only the brain tissue, we also explored if this signature is representative of the other tissues. A plausible way to approach this question is repeating the same analysis using other tissues. However, it is not straightforward because i) the number of datasets available for the other tissues limits the capacity of our approach to compile consistent signatures, increasing false positives, and ii) we find that the ageing-related changes in other tissues are not as consistent as in brain (Figure S11a). Thus, we choose another approach and asked if the direction of change for the ageing signature we compiled is similar to the direction of change in other tissues (Figure S11c). We also tested the significance of the similarity in the direction of change based on random permutations. As expected, GTEx brain data showed the highest percent similarity to the array signature. 8/35 datasets showed more dissimilarity for the down-regulated genes (i.e. percent similarity was lower than 50%), while only two were statistically significant, namely, liver and atrial appendage. Similarly, only 6/35 datasets showed more dissimilarity for the up-regulated genes, while none was significant. We repeated the analysis with the GTEx signature and observed similar results with only exception that there were five datasets with significant dissimilarity for the down-regulated genes (Figure S11e). Thus, it is possible that brain signature includes some brain-specific changes but based on significant similarity, we can say it is also representative of other tissues.

## Discussion

In this study, using gene expression data, we identified a set of drugs that are likely to modulate ageing in the human brain. Using a meta-analysis approach, we generated a reproducible ageing signature that represents multiple brain regions and is independent of the platform used for the detection of expression. Using the Connectivity Map, we identified drugs highly associated with this ageing signature. Based on the DrugAge database, seven of these drugs were previously tested on model organisms and prolonged lifespan in at least one experiment. The fact that we successfully re-discovered a statistically significant number of known lifespan modulators, without using any prior drug ageing information, suggests that the other drugs that we identified also have a high potential to be modulators of the ageing process / lifespan. Eleven of these had targets implicated in ageing, based on GenAge database (Tacutu et al., 2017). These targets include extensively studied ageing-modulators such as PI3K subunits and histone deacetylases. We also identified a group of novel candidates that are not in ageing databases, which can offer new targets and mechanisms to modulate ageing. These include drugs targeting serine / threonine, muscarinic acid, and GABA(A) receptors, protein translation, and BCL6 gene. A literature research presented in SI provides more information on potential mechanisms and suggests the list includes drugs that can influence both life- and health-span in humans.

‘Omics-based drug repurposing studies, such as the CMap, aim to identify drugs reversing the profile induced by a biological state of interest. Ageing is a time-dependent, complex phenomenon, which induces subtler changes compared to development (Dönertaş et al., 2017), or to a disease state such as Alzheimer’s (Avramopoulos, Szymanski, Wang, & Bassett, 2011). The ‘omics profile reflects two potentially distinct contributions: - the detrimental effects which occur with age (e.g. accumulation of mutations) and the potentially beneficial responses to those changes (e.g. the immune response). As a result, CMap similarity score is not conclusive on its own. In order to characterise the potential effects of drugs on ageing (anti- or pro-ageing drugs), we use three different approaches: i) comparison of the drug-induced expression profiles with the known pro-longevity drug profile (Figure S9), ii) functional analysis of the drug-induced gene expression changes (Figure S10), and iii) compilation of literature on the drugs and targets (SI). Based on these analyses we suggest that eight of seventeen drugs (quinostatin, trifluoperazine, thioridazine, vorinostat, alvespimycin, tanespimycin, rifabutin, and 15-d prostaglandin J2), which are not in DrugAge, are likely to have positive effects, whereas, topoisomerase inhibitors (camptothecin, irinotecan, and daunorubicin) can be detrimental and could act as pro-ageing drugs. Four of the remaining drugs, which are cinchonine, securinine, emetine and tretinoin, do not cluster closely with any known pro-longevity drugs in Figure S10. Literature, however, suggests cinchonine and securinine are likely to have negative effects (see SI), whereas emetine and tretinoin could act as antiageing drugs. GW-8510 and atropine oxide could not be classified because neither the clustering results nor literature evidence are conclusive.

It is important to note that none of the cell lines used to generate the CMap data originates from the brain. The assumption for using the CMap algorithm is that the effect we see in diverse cell-lines reflects the global profile of the drug perturbation and thus should be also transferable to the brain. However, it is possible that drugs have cell or tissue-specific effects. Even if the drugs induce the same expression changes in brain cells, an important question is: Can they cross the blood-brain barrier to target the brain? If some of these drugs have side effects on the CNS, it might be an indication that these drugs can affect the brain and can be re-purposed to target brain ageing. Only eight of the 24 compounds have reported side effects and all of them has at least one reported effect on the nervous system, based on MedDRA system organ classes (Table S10). This implies that these drugs can affect CNS, although we do not have information on their ability to cross the barrier. The rest may or may not cross the barrier to influence the expression in the brain, but they may also improve health by targeting generic changes throughout the body. The ageing signatures from brain tissue show a modest but significant similarity to expression profiles from non-brain tissues (Figure S10). Thus, it is possible that we identified not only drugs specifically targeting ageing in the brain but also drugs targeting ageing in other tissues. It is also possible that there are drugs which can target brain ageing with more potency, but we cannot identify them because we do not have drug-induced expression profiles for brain cells. Another important technical drawback is that the data we used to generate the ageing signature are bulk RNA expression datasets, where the expression profile is an average of all the cell types in the human brain. Focusing on the changes that are observed ubiquitously across all brain regions, we aimed to focus on global changes which are unlikely to be driven by cell type differences. However, future datasets generated using single-cell expression profiling can greatly improve the understanding of both the ageing process itself and how the interventions work.

To summarise, this study provides an unbiased identification of drugs that can target human brain ageing. We first compiled a set of gene expression changes that can characterise human brain ageing and asked if there are drugs which alter the expression of the same genes. We identified 24 drugs, seven of which were among known pro-longevity drugs. Our analysis suggests that anti-ageing drugs may act by mimicking the response while it is also possible that they can reverse the detrimental changes in ageing. Based on the literature research, we concluded that some of the drugs we identified can directly modulate the lifespan, whereas some are more likely to function by improving the cognitive functions and promoting the healthy ageing. We are in the process of experimentally testing a group of the drugs that we have identified. We hope the information presented in this study will guide research community to further test and identify chemical modulators of the ageing process in humans.

## Methods

### Data pre-processing

*Microarray datasets:* We used seven microarray-based RNA expression studies with samples from 22 brain regions, that are not mutually exclusive (Table S1). Data from different brain regions are processed and analysed separately, resulting in 26 datasets. The number of individuals in each dataset ranges between 11 and 148. The total number of individuals is 304, and the total number of samples is 805 (after removing the outliers). Some studies include samples covering the whole lifespan. However, in this study, we only considered samples above 20 years of age, which corresponds to the age at first reproduction in human societies (Walker et al., 2006). Previous human brain ageing studies using transcriptome data have also suggested gene expression patterns before and after the age of 20 are discontinuous (Colantuoni et al., 2011; Dönertaş et al., 2017). Since we are interested in finding consistent tendencies in terms of the direction of change, which can characterise ageing, we only included samples above 20 years of age. As a result, the samples included in the analysis had ages between 20-106. The microarray data were downloaded from NCBI GEO (Barrett et al., 2013) using the accession numbers in Table S1. Using “affy” (Gautier, Cope, Bolstad, & Irizarry, 2004) or “oligo” (Carvalho & Irizarry, 2010) libraries in R, RMA background correction is applied to the expression data. The data is then log2 transformed, and quantile normalized (using “preprocessCore” library in R). By visual inspection of the first and second principal components of the probe-set expression levels, outliers were excluded from the further analysis (Table S1). The age distributions for the datasets after outlier removal are given in Figure S1. Gene annotations for the probe-sets are obtained from the Ensembl database using the ‘biomaRt’ library (Durinck, Spellman, Birney, & Huber, 2009) in R. Because the annotations for the probe-sets used in Kang2011 and Colantuoni2011 are not available in Ensembl, we used the GPL files deposited in GEO. If Ensembl gene IDs are not provided in the GPL files, Entrez gene IDs were extracted and converted to Ensembl Gene IDs using the ‘biomaRt’ package. Probe-set level expression information is then mapped to gene IDs. In order not to duplicate expression values, we excluded the probe-sets corresponding to multiple genes. Expression values for the genes with multiple probe-sets were summarised using the mean expression levels. *RNA-seq dataset:* We analysed transcriptome data generated by GTEx project (v6p)(Ardlie et al., 2015). Samples are filtered based on the cause of death circumstances (4-point Hardy Scale). Only the cases with a death circumstance of 1 (violent and fast deaths due to an accident) and 2 (fast death of natural causes) are used for the downstream analysis and the samples with illnesses are excluded. Among all tissues, only the ones having at least 20 samples are considered. We also excluded ‘Cells - Transformed Fibroblasts’ category to include only the samples from tissues. As a result, 35 datasets (17 major tissue type) are used for the downstream analysis, 13 of which were from the brain. The final set that we analysed includes 2152 (623 for the brain) samples from 120 (99 for the brain) individuals. The genes with median RPKM value of 0 are also excluded from data. The RPKM values provided in the GTEx database are log2 transformed and quantile normalized for the downstream analysis. Similar to the microarray data, we excluded the outliers based on the visual inspection of the first and second principal components (Table S1). Distribution of the ages after outlier exclusion is given in Figure S1.

### Age-related expression changes and the ageing signature

The Spearman’s rank correlation coefficients between age and gene expression levels are used to measure age-related expression changes. In each dataset, we calculated the Spearman’s correlation for each gene, separately. As a result, each gene had two measures to assess its age-related expression: 1) a correlation coefficient (rho), indicating the strength and the direction of change with age and 2) a p-value, showing the significance of the association. The p-values are corrected for multiple testing using p.adjust function in R, with method=“FDR” argument. As the power to detect significant changes in each dataset is different and the sample size is small for most of the datasets, for the downstream analysis we only used the correlation coefficients (rho) and assessed the significant gene expression change tendencies that are observed in all datasets. When a gene is up-regulated by age throughout the lifespan, then it would have a positive Spearman’s correlation coefficient that is close to one. In contrast, a gene would have negative correlation coefficient if it is down-regulated. When the association is not strong, the magnitude of the correlation coefficient decreases, but the sign still reflects the direction of change that is observed in most of the time-points. We used the sign of correlation coefficient, i.e. the direction of change, to compile the set of genes that show consistent changes across all datasets. This set of genes are referred to as the ‘ageing signature’. The ageing signature, thus, does not reflect the dramatic changes in gene expression but captures consistent trends that are observed across all datasets. The statistical significance of the ageing signature is calculated using a permutation scheme, testing the significance of the consistency.

### Permutation test

We used a permutation scheme that we developed earlier (Dönertaş et al., 2017), to simulate the null hypothesis that there is no association between age and the gene expression, while retaining the dependence between genes and the datasets. Particularly, the ages of individuals in each study are permuted (randomised) 1,000 times and if that individual donated multiple samples for different brain regions, each sample is annotated with the same age. Then, the Spearman’s correlation coefficient between these randomised ages and the gene expression value for all genes are calculated. In this way, we retain the dependence between genes (e.g. those regulated by the same transcription factor) and the samples (e.g. donated by the same individuals). Permutations are performed using ‘sample’ function in base R.

Using the correlation coefficients calculated through permutations performed as explained above, we tested i) significance of the correlations among datasets, ii) significance of the finding the same or a higher number of consistently up- or down-regulated genes, i.e. the ageing signature. In order to test the significance of the correlations among datasets, we calculated the correlations between the expression-age correlation coefficients calculated using the permutations. We constructed the distribution for the median correlation coefficient among datasets (distribution of the 1,000 values), and calculated how many times the randomized values have higher correlation than the value we calculate using the real ages. In this way, we calculate an empirical p-value. The median of the permuted values reflects the value that would be expected by chance. Similarly, in order to test the significance of the ageing signature, we compiled permuted ageing signatures, for 1,000 times, and asked how many times we have the same or higher value than the calculated number of genes in the microarray or GTEx ageing signatures. In this way, we calculate the empirical p-value and median of the number of shared tendencies based on permutations, reflecting what would be expected by chance.

### Gene Ontology Enrichment

Using “topGO” and “org.Hs.eg.db” libraries in R, we performed a functional analysis of the ageing signature. Using GO categories with more than 10 annotated genes, we applied an enrichment test for the Gene Ontology (GO) (Ashburner et al., 2000) Biological Process (BP) categories.

### Connectivity Map Analysis

A list of genes showing a consistent change in ageing (the ageing signature) is used to query the Connectivity Map (Lamb, 2006). Since the Connectivity Map input requires probe-set ids, the “biomaRt” package in R is used to convert the gene list to the probeset ids that are compatible with the CMap data. The probe-sets that are in both up- and down-regulated probe-set lists are excluded from both lists. The final lists are used to query CMap database to associate the ageing signature with the drug perturbed expression profiles in the database. The resulting p-values are FDR corrected to account for multiple testing and adjusted p<0.05 is used as the significance threshold.

The ageing signature compiled using the GTEx data had more than 500 probe-sets in both up and down lists. Since the algorithm requires an input with less than 500 entries, we used the ones with the higher magnitude of expression change (median Spearman’s rank correlation coefficients across 13 brain regions). In order to show that this does not bias the results, we repeated this step for 1,000 times by randomly selecting 500 of the probe-sets in the GTEx ageing signature. In order to automatize this process, we re-implemented CMap algorithm in R and calculated the drug similarity scores using the ‘rankMatrix.txt’ data provided on the CMap website. Drug similarity scores generated using the top 500 and randomly selected 500 of the GTEx ageing signature showed a significant correlation (median rho = 0.81, range = (0.80,0.82)), suggesting that this approach does not bias the results.

### Searching the drug databases for CMap drugs

Entries in the Connectivity Map are composed of the drug names, which are generally the catalogue names for the drugs from chemical vendors. Similarly, DrugAge drugs also do not have an ID that is possible to map across different databases. The DrugAge database was retrieved on 11^th^ May 2017, from the DrugAge website. In order to compare the drugs in the Connectivity Map and the DrugAge, we first used the PubChem database (Kim et al., 2016) to make a transition across different sources. We obtained PubChem compound IDs for each drug in the Connectivity Map and DrugAge using PubChem API accessed through R programming environment and ‘RCurl’ and ‘jsonlite’ libraries.

### Targets of the drugs that are significantly associated with ageing

We compiled the drug-target associations for the drugs significantly associated with ageing mostly through literature research. For the cases where the database entries are manually curated and consistent, we used CHEMBL (Bento et al., 2014), DrugBank (Law et al., 2014), and PubChem (Kim et al., 2016). We downloaded GenAge model organism and human datasets (Tacutu et al., 2017) on 10^th^ October 2017 using GenAge website. Using the human orthologues for the model organisms (genage_models_orthologs_export.tsv) and the human dataset, we asked if any of the drug targets were previously shown to be implicated in ageing. In order to construct the drug – target network, we used ‘ggnetwork’ package in R.

### The Pro-Longevity Drug Expression Profile

In order to compile a set of gene expression changes that can be associated with the known pro-longevity drug profile, we first downloaded the pre-processed data matrix with the drug-induced expression changes (‘amplitudeMatrix.txt’ from CMap FTP server ftp://ftp.broadinstitute.org/distribution/cmap). Using this matrix, for the seven pro-longevity drugs in DrugAge that are among the significant associations according to our analysis, we generated a pro-longevity drug profile. We first identified the drug-induced gene expression changes for each of these seven drugs and each of the probe-sets that are in the microarray ageing signature. For each drug – probe-set pair, we take the direction of change that is observed in at least 60% of the experiments (using different doses or different cell lines) as the effect of that drug on the expression of that probe-set. After deciding on the individual drug effects, we took the type of change observed in at least four of seven drugs as the pro-longevity drug profile. The reason why we do not seek a perfect overlap among different drugs is to allow potentially different mechanism of actions to be included in the pro-longevity drug profile. As a result, we got five categories: 1) increase in ageing, increased by the drugs; 2) increase in ageing, decreased by the drugs; 3) decrease in ageing, increased by the drugs; 4) decrease in ageing, decreased by the drugs; and 5) the ones that are not affected consistently by the drugs. The full list of genes in the first four categories is given as TableS5. We also asked if any of the GO Biological Processes is enriched in any of the first four categories and thus did an enrichment analysis. We calculated the odds ratio for each GO category by keeping the type of change in ageing the same. For example, we asked if a GO category is enriched in genes that increase in ageing and also increased by the drugs, compared to the genes that increase in ageing but decreased by the drugs. Because the number of genes is small, it is not possible to detect significant associations after correcting for multiple testing and thus we only report the odd’s ratios for the categories (Table S6). We also compared the known prolongevity drug profile we compiled with the profile induced by the 24 drugs identified in the study (Figure S9). We calculated the percentage of probe-sets that show the same type of change as the pro-longevity drug profile. For this, we again only considered probe-sets that show the same type of change in at least 60% of the experiments per drug.

### Gene-set enrichment analysis for drug-induced changes

Using the ‘amplitudeMatrix.txt’ downloaded from the CMap website, we determined the expression changes at the gene level for each drug. We first subset the matrix to include only the experiments for the 24 significant drugs we found. We then mapped the probe-set ids (total number of probe-sets = 22,283) to Entrez gene ids using the Ensembl biomaRt package in R. We map 19,222 probe-sets to genes, excluding examples where the same probe-set id maps to multiple genes (628 multi-gene probeset ids in total). The genes with more than one probe-set id are represented by taking the median expression change induced for the probe-sets (number of genes = 12064). When the experiments for each drug are treated separately, we noticed that the results were confounded by cell-line. Thus, we then summarized multiple experiments for each drug by taking the median of the change they induce. In this way, we trimmed the cell-line specific effects. Then the expression changes (for 12064 genes) for each drug (24 drugs) are rank ordered. Using clusterProfiler package and ‘gseKEGG’ and ‘gseGO’ functions, we performed GSEA for the gene expression changes induced by each drug separately. For the KEGG pathway analysis, we only considered the pathways with at least 50 genes (188 pathways), and for GO analysis, we only considered Biological Process categories with at least 50 and maximum of 200 genes (1589 categories).

### Comparing Brain Ageing Signature to Other Tissues

We calculated the proportion of genes that show a change in the same direction with the ageing signature compiled using brain data. The proportions are calculated for ageing signatures compiled using the array and GTEx brain data, separately. We also analysed up-regulated and down-regulated genes separately to observe any differential pattern. In order to calculate the significance of similarity or dissimilarity we performed 10,000 permutations as follows: i) N number of genes, where N is the number of genes in a particular group (array / GTEx and up- / down-regulated), were selected randomly from a given GTEx dataset, ii) the proportion of changes in a given direction is calculated, and iii) using the distribution of these proportions, we asked how many times we obtain a value as extreme as the proportion calculated for that tissue and assign empirical p value.

### Side Effects

Using compound PubChem IDs, we subset the Side Effect Resource (SIDER 4.1) (Kuhn, Letunic, Jensen, & Bork, 2016), a database of adverse drugs reactions for marketed medicines. The latest version of SIDER code the side effects by using the Medical Dictionary for Regulatory Activities (MedDRA), an adverse event classification dictionary. To obtain term at the system level, we mapped the lowest-level MedDRA terms in SIDER (LLT codes) to MedDRA System Organ Class terms (SOC codes) using hierarchical files downloadable from the MedDRA web-based browser (https://tools.meddra.org/wbb/). A total of 8 drugs among the 24 had labelled side effects.

## Acknowledgments

The authors thank the whole Thornton group, especially Dobril Ivanov, Jonathan Tyzack and Daniel Elias Martin Herranz for their support and helpful discussions.

## Author Contributions

H.M.D and J.M.T designed the study. H.M.D analysed the data with the help of M.F.V.. H.M.D., J.M.T., and L.P. interpreted the results and wrote the manuscript. All authors read, revised and approved the final version of this manuscript.

## Supporting Information List

TableS1: Dataset information

TableS2: The list of genes in the ageing signatures

TableS3: The GO Enrichment results for the up-regulated genes in the microarray ageing signature

TableS4: The GO Enrichment results for the down-regulated genes in the microarray ageing signature

TableS5: The list of genes in the pro-longevity drug profile

TableS6: The GO Enrichment results for the pro-longevity drug profile

TableS7: The GO Enrichment results for the up-regulated genes in the GTEx ageing signature

TableS8: The GO Enrichment results for the down-regulated genes in the GTEx ageing signature

Table S9: Summary of the previous lifespan experiments using drugs that are re-discovered in this study.

Table S10: The list of side effects reported for the significant drugs we identified.

## References

Aliper, A., Belikov, A. V., Garazha, A., Jellen, L., Artemov, A., Suntsova, M., … Zhavoronkov, A. (2016). In search for geroprotectors: In silico screening and in vitro validation of signalome-level mimetics of young healthy state. Aging, 8(9), 2127–2152. http://doi.org/10.18632/aging.101047

Ardlie, K. G., Deluca, D. S., Segre, A. V, Sullivan, T. J., Young, T. R., Gelfand, E. T., … Dermitzakis, E. T. (2015). The Genotype-Tissue Expression (GTEx) pilot analysis: Multitissue gene regulation in humans. Science, 348(6235), 648–660. http://doi.org/10.1126/science.1262110

Ashburner, M., Ball, C. A., Blake, J. A., Botstein, D., Butler, H., Cherry, J. M., … Sherlock, G. (2000). Gene ontology: tool for the unification of biology. The Gene Ontology Consortium. Nature Genetics, 25(1), 25–29. http://doi.org/10.1038/75556

Avramopoulos, D., Szymanski, M., Wang, R., & Bassett, S. (2011). Gene expression reveals overlap between normal aging and Alzheimer’s disease genes. Neurobiology of Aging, 32(12), 2319.e27–2319.e34. http://doi.org/10.1016/j.neurobiolaging.2010.04.019

Barardo, D., Thornton, D., Thoppil, H., Walsh, M., Sharifi, S., Ferreira, S., … de Magalhães, J. P. (2017). The DrugAge database of aging-related drugs. Aging Cell, 16(3), 594–597. http://doi.org/10.1111/acel.12585

Barnes, M. R., Huxley-Jones, J., Maycox, P. R., Lennon, M., Thornber, A., Kelly, F., … De Belleroche, J. (2011). Transcription and pathway analysis of the superior temporal cortex and anterior prefrontal cortex in schizophrenia. Journal of Neuroscience Research, 89(8), 1218–1227. http://doi.org/10.1002/jnr.22647

Barrett, T., Wilhite, S. E., Ledoux, P., Evangelista, C., Kim, I. F., Tomashevsky, M., … Soboleva, A. (2013). NCBI GEO: Archive for functional genomics data sets - Update. Nucleic Acids Research, 41(D1), D991–D995. http://doi.org/10.1093/nar/gks1193

Bento, A. P., Gaulton, A., Hersey, A., Bellis, L. J., Chambers, J., Davies, M., … Overington, J. P. (2014). The ChEMBL bioactivity database: An update. Nucleic Acids Research, 42(D1), D1083–D1090. http://doi.org/10.1093/nar/gkt1031

Berchtold, N. C., Cribbs, D. H., Coleman, P. D., Rogers, J., Head, E., Kim, R., … Cotman, C. W. (2008). Gene expression changes in the course of normal brain aging are sexually dimorphic. Proceedings of the National Academy of Sciences of the United States of America, 105(40), 15605–15610. http://doi.org/10.1073/pnas.0806883105

Bjedov, I., Toivonen, J. M., Kerr, F., Slack, C., Jacobson, J., Foley, A., & Partridge, L. (2010). Mechanisms of Life Span Extension by Rapamycin in the Fruit Fly Drosophila melanogaster. Cell Metabolism, 11(1), 35–46. http://doi.org/10.1016/j.cmet.2009.11.010

Calvert, S., Tacutu, R., Sharifi, S., Teixeira, R., Ghosh, P., & de Magalhães, J. P. (2016). A network pharmacology approach reveals new candidate caloric restriction mimetics in C. elegans. Aging Cell, 15(2), 256–266. http://doi.org/10.1111/acel.12432

Carvalho, B. S., & Irizarry, R. A. (2010). A framework for oligonucleotide microarray preprocessing. Bioinformatics (Oxford, England), 26(19), 2363–2367. http://doi.org/10.1093/bioinformatics/btq431

Clancy, D. J., Gems, D., Harshman, L. G., Oldham, S., Stocker, H., Hafen, E., … Partridge, L. (2001). Extension of Life-Span by Loss of CHICO, a Drosophila Insulin Receptor Substrate Protein. Science, 292(5514), 104 LP–106. Retrieved from http://science.sciencemag.org/content/292/5514/104.abstract

Colantuoni, C., Lipska, B. K., Ye, T., Hyde, T. M., Tao, R., Leek, J. T., … Kleinman, J. E. (2011). Temporal dynamics and genetic control of transcription in the human prefrontal cortex. Nature, 478(7370), 519–523. http://doi.org/10.1038/nature10524

Dönertaş, H. M., Izgi, H., Kamacloglu, A., He, Z., Khaitovich, P., & Somel, M. (2017). Gene expression reversal toward pre-adult levels in the aging human brain and age-related loss of cellular identity. Scientific Reports, 7(1), 5894. http://doi.org/10.1038/s41598-017-05927-4

Duran-Frigola, M., Mateo, L., & Aloy, P. (2017). Drug repositioning beyond the low-hanging fruits. Current Opinion in Systems Biology, 3, 95–102. http://doi.org/10.1016/j.coisb.2017.04.010

Durinck, S., Spellman, P. T., Birney, E., & Huber, W. (2009). Mapping identifiers for the integration of genomic datasets with the R/Bioconductor package biomaRt. Nature Protocols, 4(8), 1184–1191. http://doi.org/10.1038/nprot.2009.97

Fontana, L., & Partridge, L. (2015). Promoting health and longevity through diet: From model organisms to humans. Cell, 161(1), 106–118. http://doi.org/10.1016/j.cell.2015.02.020

Fontana, L., Partridge, L., & Longo, V. D. (2010). Extending healthy life span--from yeast to humans. Science (New York, N.Y.), 328(5976), 321–326. http://doi.org/10.1126/science.1172539

Gautier, L., Cope, L., Bolstad, B. M., & Irizarry, R. A. (2004). affy--analysis of Affymetrix GeneChip data at the probe level. Bioinformatics (Oxford, England), 20(3), 307–315. http://doi.org/10.1093/bioinformatics/btg405

Kang, H. J., Kawasawa, Y. I., Cheng, F., Zhu, Y., Xu, X., Li, M., … Sestan, N. (2011). Spatio-temporal transcriptome of the human brain. Nature, 478(7370), 483–489. http://doi.org/10.1038/nature10523

Kim, S., Thiessen, P. A., Bolton, E. E., Chen, J., Fu, G., Gindulyte, A., … Bryant, S. H. (2016). PubChem substance and compound databases. Nucleic Acids Research, 44(D1), D1202–D1213. http://doi.org/10.1093/nar/gkv951

Kuhn, M., Letunic, I., Jensen, L. J., & Bork, P. (2016). The SIDER database of drugs and side effects. Nucleic Acids Research, 44(D1), D1075–D1079. http://doi.org/10.1093/nar/gkv1075

Lamb, J. (2006). The Connectivity Map: Using Gene-Expression Signatures to Connect Small Molecules, Genes, and Disease. Science, 313(5795), 1929–1935. http://doi.org/10.1126/science.1132939

Law, V., Knox, C., Djoumbou, Y., Jewison, T., Guo, A. C., Liu, Y., … Wishart, D. S. (2014). DrugBank 4.0: Shedding new light on drug metabolism. Nucleic Acids Research, 42(D1), D1091–D1097. http://doi.org/10.1093/nar/gkt1068

Lu, T., Pan, Y., Kao, S.-Y., Li, C., Kohane, I., Chan, J., & Yankner, B. A. (2004). Gene regulation and DNA damage in the ageing human brain. Nature, 429(6994), 883–891. http://doi.org/10.1038/nature02661

Lucanic, M., Lithgow, G. J., & Alavez, S. (2013). Pharmacological lifespan extension of invertebrates. Ageing Research Reviews. http://doi.org/10.1016/j.arr.2012.06.006

Mair, W., & Dillin, A. (2008). Aging and Survival: The Genetics of Life Span Extension by Dietary Restriction. Annual Review of Biochemistry, 77(1), 727–754. http://doi.org/10.1146/annurev.biochem.77.061206.171059

Mannick, J. B., Del Giudice, G., Lattanzi, M., Valiante, N. M., Praestgaard, J., Huang, B., … Klickstein, L. B. (2014). mTOR inhibition improves immune function in the elderly. Science Translational Medicine, 6(268), 268ra179. http://doi.org/10.1126/scitranslmed.3009892

Maycox, P. R., Kelly, F., Taylor, A., Bates, S., Reid, J., Logendra, R., … de Belleroche, J. (2009). Analysis of gene expression in two large schizophrenia cohorts identifies multiple changes associated with nerve terminal function. Molecular Psychiatry, 14(12), 1083–1094. http://doi.org/10.1038/mp.2009.18

Moskalev, A., Chernyagina, E., de Magalhães, J. P., Barardo, D., Thoppil, H., Shaposhnikov, M., … Zhavoronkov, A. (2015). Geroprotectors.org: A new, structured and curated database of current therapeutic interventions in aging and age-related disease. Aging, 7(9), 616–628. http://doi.org/10.18632/aging.100799

Naumova, O. Y., Palejev, D., Vlasova, N. V, Lee, M., Rychkov, S. Y., Babich, O. N., … Grigorenko, E. L. (2012). Age-related changes of gene expression in the neocortex: preliminary data on RNA-Seq of the transcriptome in three functionally distinct cortical areas. Development and Psychopathology, 24(4), 1427–1442. http://doi.org/10.1017/S0954579412000818

Niccoli, T., & Partridge, L. (2012). Ageing as a Risk Factor for Disease. Current Biology, 22(17), R741–R752. http://doi.org/10.1016/j.cub.2012.07.024

Pearson, K. J., Baur, J. A., Lewis, K. N., Peshkin, L., Price, N. L., Labinskyy, N., … de Cabo, R. (2008). Resveratrol Delays Age-Related Deterioration and Mimics Transcriptional Aspects of Dietary Restriction without Extending Life Span. Cell Metabolism, 8(2), 157–168. http://doi.org/10.1016/j.cmet.2008.06.011

Somel, M., Guo, S., Fu, N., Yan, Z., Hu, H. Y., Xu, Y., … Khaitovich, P. (2010). MicroRNA, mRNA, and protein expression link development and aging in human and macaque brain. Genome Research, 20(9), 1207–1218. http://doi.org/10.1101/gr.106849.110

Somel, M., Khaitovich, P., Bahn, S., Pääbo, S., & Lachmann, M. (2006). Gene expression becomes heterogeneous with age. Current Biology?: CB, 16(10), R359–60. http://doi.org/10.1016/j.cub.2006.04.024

Somel, M., Liu, X., Tang, L., Yan, Z., Hu, H., Guo, S., … Khaitovich, P. (2011). MicroRNA-driven developmental remodeling in the brain distinguishes humans from other primates. PLoS Biology, 9(12), e1001214. http://doi.org/10.1371/journal.pbio.1001214

Strong, R., Miller, R. A., Astle, C. M., Baur, J. A., De Cabo, R., Fernandez, E., … Harrison, D. E. (2013). Evaluation of resveratrol, green tea extract, curcumin, oxaloacetic acid, and medium-chain triglyceride oil on life span of genetically heterogeneous mice. Journals of Gerontology - Series A Biological Sciences and Medical Sciences, 68(1), 6–16. http://doi.org/10.1093/gerona/gls070

Tacutu, R., Thornton, D., Johnson, E., Budovsky, A., Barardo, D., Craig, T., … de Magalhaes, J. P. (2017). Human Ageing Genomic Resources: 2018 Update. Doi.Org, 193326. http://doi.org/10.1101/193326

Walker, R., Gurven, M., Hill, K., Migliano, A., Chagnon, N., De Souza, R., … Yamauchi, T. (2006). Growth rates and life histories in twenty-two small-scale societies. American Journal of Human Biology, 18(3), 295–311. http://doi.org/10.1002/ajhb.20510

Xiao, R., Zhang, B., Dong, Y., Gong, J., Xu, T., Liu, J., & Xu, X. Z. Z. S. (2013). A genetic program promotes C. elegans longevity at cold temperatures via a thermosensitive TRP channel. Cell, 152(4), 806–817. http://doi.org/10.1016/j.cell.2013.01.020

Xue, H., Xian, B., Dong, D., Xia, K., Zhu, S., Zhang, Z., … Han, J.-D. J. (2007). A modular network model of aging. Molecular Systems Biology, 3(1), 147. http://doi.org/10.1038/msb4100189

Ye, X., Linton, J. M., Schork, N. J., Buck, L. B., & Petrascheck, M. (2014). A pharmacological network for lifespan extension in Caenorhabditis elegans. Aging Cell, 13(2), 206–215. http://doi.org/10.1111/acel.12163

Zhavoronkov, A., Buzdin, A. A., Garazha, A. V., Borisov, N. M., & Moskalev, A. A. (2014). Signaling pathway cloud regulation for in silico screening and ranking of the potential geroprotective drugs. Frontiers in Genetics, 5(MAR). http://doi.org/10.3389/fgene.2014.00049

Ziehm, M., Kaur, S., Ivanov, D. K., Ballester, P. J., Marcus, D., Partridge, L., & Thornton, J. M. (2017). Drug repurposing for aging research using model organisms. Aging Cell, 16(5), 1006–1015. http://doi.org/10.1111/acel.12626

